# Rare Variants in the *OTOG* Gene Are a Frequent Cause of Familial Meniere’s Disease

**DOI:** 10.1101/771527

**Authors:** Pablo Roman-Naranjo, Alvaro Gallego-Martinez, Andrés Soto-Varela, Ismael Aran, Maria del Carmen Moleon, Juan Manuel Espinosa-Sanchez, Juan Carlos Amor-Dorado, Angel Batuecas-Caletrio, Paz Perez-Vazquez, Jose A. Lopez-Escamez

**Author notes:** All correspondence should be addressed to: José Antonio López Escámez, Otology & Neurotology Group CTS495, GENYO, - Centre for Genomics and Oncological Research-Pfizer/Universidad de Granada/Andalusian Regional Government, Avda de la Ilustración 114, Granada 18016 Spain.

## Abstract

**Objectives:** Meniere’s disease (MD) is a rare inner ear disorder characterized by sensorineural hearing loss, episodic vertigo and tinnitus. Familial MD has been reported in 6-9% of sporadic cases, and few genes including *FAM136A, DTNA, PRKCB, SEMA3D* and *DPT* have been involved in single families, suggesting genetic heterogeneity. In this study, the authors recruited 46 families with MD to search for relevant candidate genes for hearing loss in familial MD.

**Design:** Exome sequencing data from MD patients were analyzed to search for rare variants in hearing loss genes in a case-control study. A total of 109 patients with MD (73 familial cases and 36 early-onset sporadic patients) diagnosed according to the diagnostic criteria defined by the Barany Society were recruited in 11 hospitals. The allelic frequencies of rare variants in hearing loss genes were calculated in individuals with familial MD. A single rare variant analysis (SRVA) and a gene burden analysis (GBA) were conducted in the dataset selecting one patient from each family. Allelic frequencies from European and Spanish reference datasets were used as controls.

**Results:** A total of 5136 single nucleotide variants in hearing loss genes were considered for SRVA in familial MD cases, but only one heterozygous variant in the *OTOG* gene (rs552304627) was found in two unrelated families. The GBA found an enrichment of rare missense variants in the *OTOG* gene in familial MD. So, 15/46 families (33%) showed at least one rare missense variant in the *OTOG* gene, suggesting a key role in familial MD.

**Conclusions:** The authors found an enrichment of multiplex rare missense variants in the *OTOG* gene in familial MD. This finding supports *OTOG* as a relevant gene in familial MD and set the groundwork for genetic testing in MD.

## INTRODUCTION

Meniere’s disease [MD (OMIM 18600)] is a rare inner ear disorder with three major symptoms: sensorineural hearing loss (SNHL), episodic vertigo and tinnitus (Lopez-Escamez et al. 2015; Espinosa-Sanchez & Lopez-Escamez 2016). Hearing loss always involves low and medium frequencies in one or both ears (unilateral or bilateral MD) at the onset of the disease. However, MD also affects high frequencies in early or advanced stages of the disease (Belinchon et al. 2011). Epidemiological studies indicate that MD is most common in European population, suggesting a genetic predisposition (Ohmen et al. 2013). Although the majority of MD patients are considered sporadic (Frejo et al. 2016; Frejo et al. 2017), familial clustering has been reported in 8-9% of sporadic cases in the European descendent (Requena et al. 2014), and in 6% of Korean population (Lee et al. 2015), which also supports a genetic contribution to the disease (Roman-Naranjo et al. 2017). MD shows a wide range of phenotypic variations among patients, even within the same families (Lee et al. 2015b), and it is commonly associated with migraine and systemic autoimmune disorders (Tyrrell et al. 2014; Cha et al. 2008). Familial MD (FMD) shows an autosomal dominant (AD) pattern of inheritance with incomplete penetrance and anticipation, showing an earlier onset compared to sporadic cases (Morrison et al. 2009; Birgerson et al. 1987; Klar et al. 2006). Different whole exome sequencing (WES) based studies have identified several genes related with FMD. Single nucleotide variants (SNV) in *DTNA*, *FAM136A*, *PRKCB*, *DPT* and *SEMA3D* were identified in 4 different families AD inheritance with incomplete penetrance (Requena et al. 2015; Martín-Sierra et al. 2016; Martín-Sierra et al. 2017). However, these findings have not been replicated neither in other MD families nor sporadic MD (SMD) cases.

WES continues to be an efficient tool to determine disease-causing variants (Williams et al. 2016; Adams & Eng 2018; Suwinski et al. 2019), although the monogenic hypothesis in FMD should be reconsidered to achieve results beyond private rare variants for singular families. Thus, the “one variant-one disease” hypothesis, described for classic Mendelian inheritance cannot explain the incomplete penetrance or variable expressivity observed in MD (Martín-Sierra et al. 2017) and more complex inheritance models are needed (Cooper et al. 2013; Kousi & Katsanis 2015). Oligogenic and multiallelic models have been already applied in different diseases, such as Parkinson (Lubbe et al. 2016) and Huntington’s disease (Lee et al. 2015a), explaining changes in disease progression and phenotypic variability. Furthermore, a digenic inheritance of deafness was reported by variants in *CDH23* and *PCDH15* (Zheng et al. 2005), and recently, an enrichment of rare missense variants in certain SNHL genes, such as *GJB2*, *SLC26A4* or *USH1G*, was found in a large cohort of SMD cases (Gallego-Martinez et al. 2019), supporting the hypothesis of multiallelic inheritance in MD.

More than 150 genes have been associated to deafness (Azaiez et al. 2018), and 116 of them are related with non-syndromic SNHL (Van Camp G. 2018).

In this study, we have investigated the genetic background of FMD, focusing on SNHL genes by analyzing 46 families with MD by WES. We have found an enrichment of rare missense variants in the *OTOG* gene compared with non-Finnish European (NFE) and Spanish populations. The *OTOG* gene, which encodes otogelin, has been previously associated with deafness and imbalance and causes autosomal recessive deafness 18B (Simmler et al. 2000a; Schraders et al. 2012). A total of 15 families out of 46 showed, at least, one rare missense variant in this gene, suggesting a key role of otogelin in MD.

## MATERIALS AND METHODS

### Patient assessment and selection

A total of 73 MD patients from 46 different families with one or more affected first-degree relatives, and 36 sporadic MD cases with an age of onset younger than 35 were recruited. Patients were diagnosed following the diagnostic criteria described by the International Classification Committee for Vestibular Disorders of the Barany Society (Lopez-Escamez et al. 2015). A complete hearing and vestibular assessment was carried out in all cases, including a brain magnetic resonance imaging to exclude other causes of neurological symptoms. Serial pure tone audiograms were retrieved from clinical records to assess hearing loss since the initial diagnosis. A summary of the clinical information of these patients is presented in the Supplemental Digital Content 1 (see Table 1 to Table 3, Supplemental Digital Content 1).

**Table 1.** Rare Variants Found in the Gene Burden Analysis in *OTOG* Gene for Familial MD Cases.

| Variant position | Exon | Families | Sporadic cases | MAF FMD | MAF ALL MD | MAF NFE | MAF CSVS | CADD | Domain |
| --- | --- | --- | --- | --- | --- | --- | --- | --- | --- |
| 11:17574758G>A | 4 | F1; F14 | – | 0.041 (3/73) | 0.028 (3/109) | 0.00080 | 0.0033 | 24.8 | vWD |
| 11:17578774G>A | 7 | F2; F3; F4; F5 | S24 | 0.068 (5/73) | 0.055 (6/109) | 0.0090 | 0.017 | 15.95 | vWD |
| 11:17594747C>A* | 18 | F34 | – | 0.027 (2/73) | 0.018 (2/109) | – | – | 22.2 | C8 |
| 11:17621218C>T | 30 | F6; F7 | – | 0.027 (2/73) | 0.018 (2/109) | 0.0026 | 0.0033 | 34 | C8 |
| 11:17627548G>A | 32 | F14 | – | 0.012 (1/73) | 0.009 (1/109) | 0.0056 | 0.0054 | 23.6 | Abf |
| 11:17631453C>T | 35 | F8 | S11; S24 | 0.012 (1/73) | 0.028 (3/109) | 0.017 | 0.014 | 12.89 | – |
| 11:17632921C>T | 35 | F2; F3; F4; F5 | – | 0.068 (5/73) | 0.046 (5/109) | 0.0015 | 0.0054 | 7.71 | – |
| 11:17656672G>A | 45 | F10 | S9 | 0.013 (1/73) | 0.018 (2/109) | 0.0034 | 0.0039 | 31 | – |
| 11:17663747G>A | 52 | F1; F13; F14 | S7 | 0.055 (4/73) | 0.046 (5/109) | 0.0058 | 0.0054 | 19.41 | – |
| 11:17667139G>C | 54 | F9; F11; F12 | S12; S20 | 0.082 (6/73) | 0.073 (8/109) | 0.019 | 0.017 | 27.2 | CT |

\* This novel variant was not included in the gene burden analysis. Abbreviations: MAF FMD, minor allele frequency in familial MD; MAF ALL MD, minor allele frequency in all familial and non-familial MD cases ; NFE, Minor allele frequency in non-Finnish European population; CSVS, Collaborative Spanish Variant Server; Abf, alpha-L-arabinofuranosidase B domain; CADD, Combined Annotation Dependent Depletion Score; CT: cysteine knot domain; vWD, von Willebrand factor type D domain.

This study protocol was approved by the Institutional Review Board for Clinical Research (MS/2014/02), and a written informed consent to donate biological samples was obtained from all subjects.

### DNA extraction and whole exome sequencing

Blood and saliva samples were taken from patients with MD to perform WES. DNA samples were extracted with prepIT-L2P (DNA Genotek, Ottawa, Canada) and QIAamp DNA Blood Mini Kit (Qiagen, Venlo, The Netherlands) using manufacturer’s protocols and quality controls previously described (Szczepek et al. 2019). DNA libraries were prepared by using the SureSelect Human All Exon V6 kit (Agilent Technologies, Santa Clara, CA, USA) and were paired-end sequenced on the Illumina HiSeq 4000 platform at 100X coverage. Raw reads were stored in two FASTQ files for each individual.

### Bioinformatic analysis

#### Dataset generation and processing

Analysis-ready BAM files and VCF files were generated from raw unmapped reads using the GATK Best Practices pipeline. Reads were aligned to the GRCh37/hg19 human reference genome using the BWA-MEM algorithm. For obtaining the final dataset, SNV and small structural variants were filtered according to its Variant Quality Score Recalibration (VQSR) and depth of coverage (DP) values. Thus, variants were excluded if their VQSR value were under the VSQR threshold or their average DP < 10. Variants were functionally annotated using ANNOVAR version 2018Apr16. RefSeq was used for gene-based annotation and the Exome Aggregation Consortium (ExAC) database, the Combined Annotation Dependent Depletion (CADD) scores and the dbNSFP database (v3.0) were used for filter-based annotation.

#### Sensorineural hearing loss gene set

The SNHL gene set was generated by using three different databases: the Hereditary Hearing Loss Homepage (Van Camp G. 2018), the Deafness Variation Database (Azaiez et al. 2018) and Harmonizome (Rouillard et al. 2016), containing a total of 116 genes related with SNHL (see Table 4, Supplemental Digital Content 1).

#### Data analysis and prioritization strategy

Two pipelines and filtering/prioritization strategies were conducted to search for rare variants as we have previously described (Gallego-Martinez et al. 2019). The first was a single rare variant analysis (SRVA) for studying individual families; the second approach was a gene burden analysis (GBA) to obtain a gene-level mutational profile (Figure 1). For these analyses only one patient from each family was selected.

**Figure 1:**
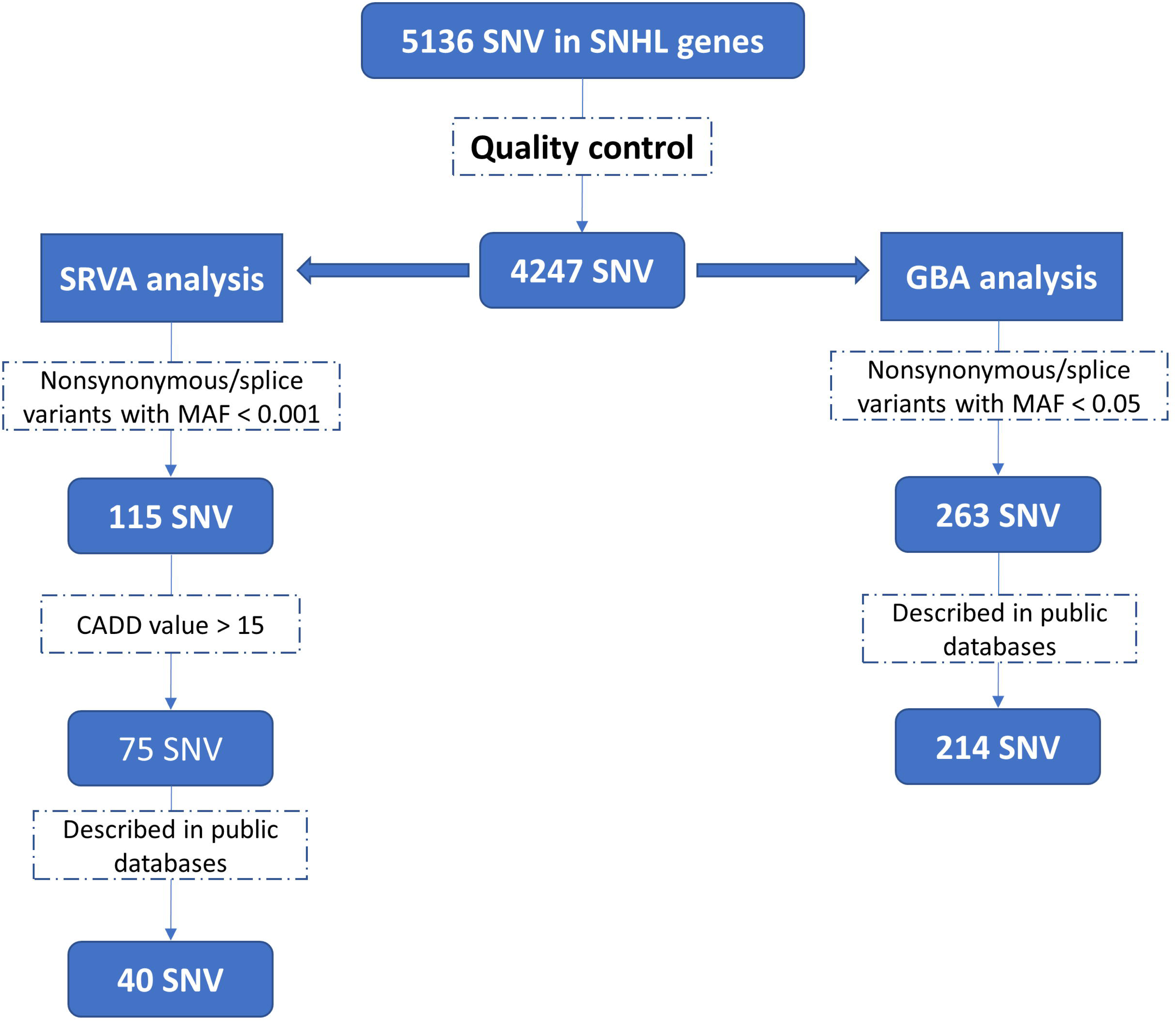
Flowchart summarizing the bioinformatic analysis on familial MD cases. On the left, single rare variant analysis (SRVA) and prioritization pipeline. On the right, the gene burden analysis (GBA) pipeline. SNV, single nucleotide variants; CADD, Combined Annotation Dependent Depletion Score.

Whenever possible, the patient selected was in the last generation. Sporadic cases with an early onset were also investigated to search for singleton variants in candidate genes in both analyses.

All variants were assessed according to the standards and guidelines described by the American College of Medical Genetics and Genomics (ACMG) and the Association for Molecular Pathology (AMP) (Richards et al. 2015). Variants not described in the NFE population from ExAC and the Spanish population from CSVS were discarded to minimize false calls and population-specific variants (Shearer et al. 2014). Selected variants were checked in patients BAM files with IGV and/or sequenced by Sanger sequencing to minimize false calls.

#### Statistics and databases

Two independent datasets were used as reference to compare the observed MAF in FMD and to calculate odds ratios (OR): NFE population from ExAC and the Collaborative Spanish Variant Server (CSVS) database (Lek et al. 2016; Dopazo et al. 2016).

For each selected variant in the SRVA, OR with 95% confidence interval (CI) were calculated using the MAF values from the CSVS database (N=1,579) and the NFE population (N=33,365) from ExAC.

For GBA, we counted the total exonic alternate alleles per gene in our cohort against the two reference datasets. After calculating OR with 95% CI, we obtained one-sided p-values that were corrected for multiple testing by the total number of variants found in each gene following the Bonferroni approach.

Standard audiometric evaluations for air and bone conduction elicited by pure tones from 125 to 8000 Hz were retrieved from the clinical records to analyse the time course of the hearing profile in FMD cases with candidate variants. Regression analysis was performed to estimate the outcome of hearing loss for each frequency.

## RESULTS

### Main genetic findings in familial MD

#### Single rare variant analysis

A total of 5136 variants located in SNHL genes were considered in FMD cases. After applying quality controls (QC), 4247 SNV remained. Only 75 nonsynonymous or splice site SNV fulfilled the MAF (<0.001) and CADD (>15) filtering criteria (Figure 1).

From them, 40 SNV were already described in the NFE population or Spanish population, but only one SNV was found in more than one family (see Table 5, Supplemental Digital Content 1, which shows the rare variants found in the SRVA for FMD cases). This heterozygous variant located in *OTOG* gene was observed in cases from two unrelated families (F1 & F14). The variant chr11:17574758G>A (rs552304627; p.V141M), which is in the last nucleotide of the fourth exon in the *OTOG* canonical transcript (ENST00000399391), is likely pathogenic according to the ACMG and AMP guidelines. This multiplex variant is located in a Von Willebrand Factor D-type domain (vWD) with a MAF=.0008 in NFE population, and multiple *in silico* tools supported a likely pathogenic effect of this variant (SIFT score=.001; M-CAP=.153; CADD=28.2; GERP++ =5.36).

The rest of the rare SNV were considered private familial variants because none of them were found in other FMD cases. None small structural variant (insertion or deletion) was found in any SNHL genes.

#### Gene burden analysis

Seventy-three genes with 214 SNV with a MAF<0.05 were retained after QC and filtering steps. Most of the genes (74%) carried less than 3 variants, thus they were discarded for further analysis. The most significant finding was an enrichment of rare missense variants in *OTOG* gene in our FMD cases against either NFE population from ExAC (OR= 4.3(2.6-7.0), *p*= 4.1×10^-8^) or Spanish population (OR= 3.6(2.1-5.9), *p*= 7.1×10^-6^). Nine different rare missense variants were found in *OTOG* in 14/46 non-related families, existing 6 families with 2 or more shared variants (Table 1 & see Figure 1, Supplemental Digital Content 2). The variants rs61978648 and rs61736002 were shared by individuals from 4 unrelated families (F2, F3, F4 & F5). Likewise, the variants rs552304627 and rs117315845 were found in patients from other 2 unrelated families (F1 & F14).

In addition, a novel variant in *OTOG* not included in the GBA was found in two cases from a 15th family (F34). This variant, located in exon 18 (chr11:17594747C>A), was found in heterozygous state affecting the sequence of the C8 domain. The distribution of the variants found in *OTOG* is scattered across the gene sequence (Figure 2).

**Figure 2:**
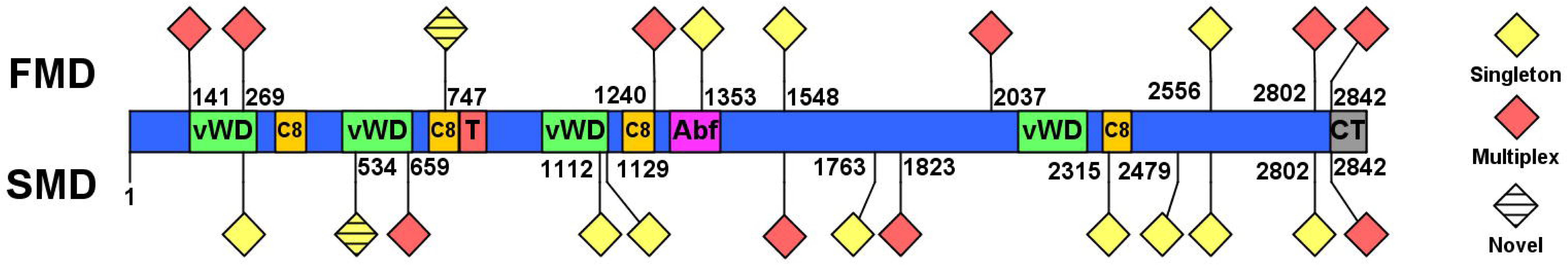
Variants distribution across OTOG gene domains. On the upper part, variants which were found in familial Meniere disease (FMD) cases. On the bottom part, variants which were found in sporadic Meniere disease (SMD) cases. Yellow-colored variants indicate variants found in only one case, whereas red-colored variants represent variants found in 2 or more cases in a cohort. vWD, von Willebrand factor type D domain; T, Trypsin inhibitor-like domain; Abf, Alpha-L-arabinofuranosidase B domain; CT, Cysteine knot domain.

### Hearing profile in familial patients with rare variants in *OTOG*

The hearing profile for the 14 patients (3 males, 11 females) with rare variants in *OTOG* gene was studied (see Figure 2, Supplemental Digital Content 2, which shows the pure tone audiograms for these patients). Ten of them showed bilateral hearing loss, 3 had left-sided hearing loss and only 1 patient shown right-sided SNHL (see Table 1, Supplemental Digital Content 1, which summarizes the clinical information of the familial MD cases carrying variants in *OTOG* gene). From these 14 patients, 16 ears from 12 patients showed a flat shaped audiogram (57.1%), 5 ears from 5 patients showed a ski-slope shaped audiogram (17.8%), 3 ears from 3 patients showed a reverse-slope shaped (10.7%) and 4 ears had a normal pure-tone audiogram (14.2%).

A regression analysis was done to estimate the hearing loss at onset and the outcome for each frequency. We found a negative correlation at 1000 Hz (R^2^=.143; *p*=.033) and 2000 Hz (R^2^=.246; *p*=.004). There was no statistical correlation at 125 Hz, 250 Hz, 500 Hz, 4000 Hz nor 8000 Hz, suggesting no progression at these frequencies (Figure 3). The age of onset of the symptoms was 41.93±8.66 and the estimated hearing loss at onset was 62.14±12.83 for low frequencies (125-250-500 Hz) and 58.75±14.1 for high frequencies (1000-2000-4000 Hz).

**Figure 3:**
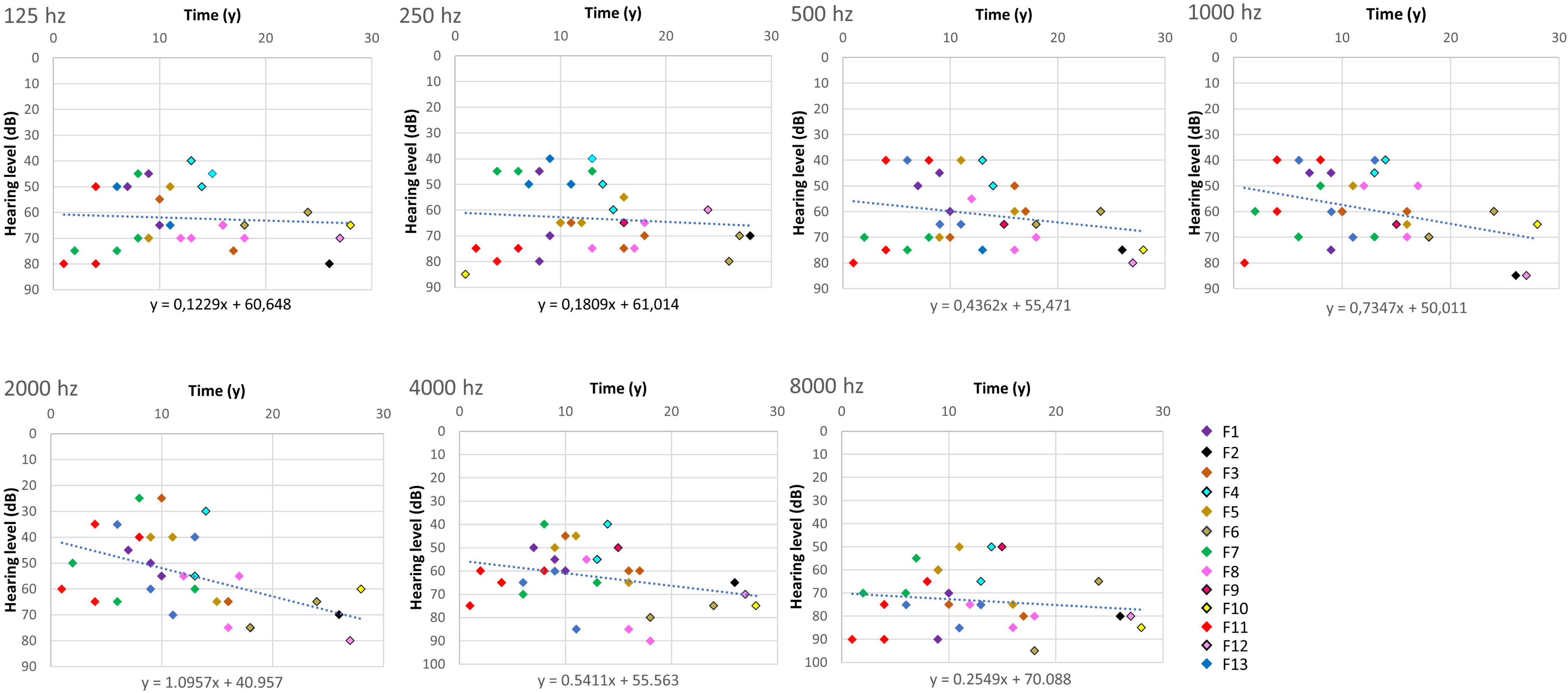
Scattered plot showing air conduction hearing thresholds obtained and the duration of the disease for each frequency in familial MD cases. Regression equations and estimated hearing loss at the onset are displayed below the charts.

### Early onset sporadic MD

The same analytical pipeline was used in a series of patients with sporadic MD with an age of onset younger than 35 (see Figure 3, Supplemental Digital Content 2). For the SRVA, we found 60 nonsynonymous or splice site SNV with MAF < 0.001 and CADD > 15 in SNHL genes. Among them, one variant was found in two sporadic cases and another variant was also found in a familial case. The rest of the SNV were considered simplex variants found in singletons and none of them were homozygous (see Table 6, Supplemental Digital Content 1, which shows the rare variants found in the SRVA for SMD cases).

A heterozygous nonsynonymous SNV in *OTOG* gene was found in two unrelated sporadic MD cases (S1 and S23). The variant chr11:17632279C>T (rs779658224; p.A1823V) is located in exon 35 of the canonical transcript of *OTOG* gene and it is a variant of uncertain significance (VUS) according to the ACMG and AMP guidelines. This variant has a MAF=.0005 in the NFE population from ExAC and it is not described in the Spanish population from the CSVS. In addition, a heterozygous nonsynonymous SNV in *OTOGL* gene was found in one sporadic case and in one familial case (S27 and F31). The variant chr12:80752642T>G (rs145929269; p.C2068G) is located in exon 51 of the canonical transcript of *OTOGL* gene (ENST00000458043). This region encodes a cysteine-rich region and this variant was also classified as a VUS according to the ACMG and AMP guidelines.

For the GBA, we found 12 rare SNV in *OTOG* gene in patients with early onset MD (see Table 7, Supplemental Digital Content 1). However, in contrast with the results obtained in FMD cases, there was not an excess of rare variants in this gene against neither the NFE population from ExAC (OR=2.1(1.2-3.7), *p*=.11) nor Spanish population (OR=2(1.1-3.5), *p*=.20) (Figure 2).

## DISCUSSION

Familial MD has an AD inheritance with incomplete penetrance (Morrison et al. 2009; Requena et al. 2014), and few genes have been involved in singular families (Requena et al. 2015; Martín-Sierra et al. 2016; Martín-Sierra et al. 2017). In this study, we have found an enrichment of rare missense variants in several unrelated patients with FMD in the *OTOG* gene. These variants were observed in 15 of 46 non-related families (33% familial cases). Seven of the 15 families with rare variants in *OTOG* showed incomplete penetrance (47%) and partial syndromes (episodic vertigo or hearing loss) were found in relatives from 5 of 15 families (Morrison et al. 2009; Requena et al. 2015; Martín-Sierra et al. 2016; Martín-Sierra et al. 2017). Most of these rare variants were found in 2, 3 or 4 unrelated individuals from different families with MD and they were considered multiplex variants. However, the majority of the variants in *OTOG* found in non-familial patients with early onset were not observed in other sporadic cases (singletons variants).

*OTOG*, which encodes otogelin, was described for first time by Cohen-Salmon *et al* (Cohen-Salmon et al. 1997). Otogelin is a 2925 amino acid protein (ENST00000399391) constituted by several vWD and C8 domains, and a cysteine knot-like domain in its C-terminal. It is mainly expressed in acellular structures which cover the sensory inner ear epithelia: the tectorial membrane, the otoconial membranes and the cupula over the cristae ampullaris of the semicircular canals. Because of its localization in the extracellular structures overlying the stereocilia of the hair cells involved in the mechanotransduction of sound and acceleration, this structural protein plays an important role in both auditory and vestibular functions (Schrauwen et al. 2016).

The effects of variants in otogelin were first demonstrated in the orthologous gene in a mouse model. Three mouse models have been generated for evaluating the phenotypic changes resulting from *OTOG* variants. In the Otog^tm1Prs^ model, authors inactivated Otog by deleting the first three exons. Vestibular dysfunction was detected at P4 in Otog^-/-^, observing anomalies in the saccule and utricule. The auditory function was evaluated by Pleyer reflex, showing profound hearing impairment. The Otog^+/−^ mice did not present any anomalies (Simmler et al. 2000a). The second model is the twister (twt) mice, mice with a spontaneous recessive mutation entailing absence of Otog expression. Similarly to Otog^tm1Prs^, in Otog^twt^ the vestibular dysfunction was detected at P4, and the hearing loss was progressive and moderate to severe/profound (Simmler et al. 2000b). The last mouse model published is the otogelin ENU-induced mouse model. In this model, a homozygous variant at the splice donor site of intron 29, Otog^vbd/vbd^, cause a frame-shift and a premature codon. Otog^vbd/vbd^ mice shown abnormal hearing and vestibular functions (El Hakam Kamareddin et al. 2015).

Four variants have been described in *OTOG* gene causing DFNB18B. Schraders *et al*. were the first to describe causative variants in *OTOG*. A homozygous 1bp deletion, c.5508delC (p.Ala1838Profs*31) in four related patients, and two compound-heterozygous variants, c.6347C>T (p.Pro2116Leu) and c.6559C>T (p.Arg2187*) in other two related patients, were described to cause hearing loss and vestibular dysfunction (Schraders et al. 2012). More recently, a homozygous nonsense variant c.330C>G (p.Tyr110*) in a Korean patient has been identified, showing early-onset mild hearing loss without vestibular dysfunction (Yu et al. 2019).

In contrast to studies mentioned above, *OTOG* variants found in this study were all in heterozygous state and, despite 6 FMD cases and 3 SMD cases studied had two or more variants, compound heterozygous variants could not be demonstrated because samples from the parents were not available and *OTOG* variant segregation could not be fully assessed in each family. However, the variants chr11:17574758G>A and chr11:17663747G>A found in F14 were also identified in his mother, the variants chr11:17578774G>A and chr11:17632921C>T found in F5 were also identified in her sister (II-7), and a novel variant chr11:17594747C>A not considered for the GBA were found in F34 and her brother. Furthermore, variants located in untranslated regions (UTRs) and promoter regions, which modulate gene expression and different protein features (Chatterjee & Pal 2009; Buckland 2006), could not be evaluated because of the study design. Altogether, the results obtained by GBA suggested a different genetic architecture in FMD cases and SMD cases, since the enrichment of rare variants in *OTOG* gene was only found in FMD cases and most of the variants found in sporadic cases with early onset were singletons (not observed in multiple individuals).

Each region of the cochlea is specifically stimulated by a specific frequency. Thus, the base of the cochlea mainly responds to high-frequency sounds, whereas the apex responds to low-frequency sounds, frequencies mostly affected in MD (Robles & Ruggero 2017; Nakashima et al. 2016). Of note, otogelin shows a tonotopic gene expression in mice (Yoshimura et al. 2014). *OTOG* gene showed a 2.43-fold change in expression for apex vs base, making this gene a possible candidate for SNHL in MD. In addition, an RNA-seq study of the inner ear from patients with normal hearing showed a high expression of *OTOG* gene in the vestibule (Schrauwen et al. 2016), which could explain the vestibular dysfunction in patients with pathogenic variants in this gene.

The audiograms of FMD patients who carried rare variants in *OTOG* gene showed a moderate-to-severe flat hearing loss ≈60 dB since the first years of onset involving all frequencies. Low-frequency hearing had slight variations throughout the years, while a negative correlation was found at mid (1000Hz) and high-frequency (2000Hz) hearing.

Data from F14 were considered as an outlier and discarded because his hearing profile was not comparable to the rest of FMD patients (see Figure 2, Supplemental Digital Content 2). Since all frequencies are involved since the onset of the disease, we can speculate that the damage of the tectorial membrane mediated by mutations in otogelin will involve the entire cochlea from base to apex.

According to our results, the clinical picture of patients with mutations in *OTOG* would be a female of 43 years old with sudden or rapidly progressive flat SNHL around 60 dB and vertigo attacks with a family history of MD, vertigo or early onset SNHL.

In conclusion, we have found an enrichment of rare missense variants in the *OTOG* gene in FMD cases. These findings support a multiallelic contribution in MD, where *OTOG* gene seems to be playing a relevant role in the pathophysiology of hearing and vestibular functions in MD.

## Supporting information

Supplemental Digital Content 1

Supplemental Digital Content 1

## Acknowledgements

We thank to all participants of the Meniere’s Disease Consortium for recruiting patients with familial MD and their relatives. Pablo Roman-Naranjo is a PhD student in the Biomedicine Program at Universidad de Granada and his salary was supported by ASMES (Asociación Sindrome de Meniere España). JALE conceived the study design and recruited all clinicians involved in the Meniere’s Disease Consortium to characterize families with MD at different sites (AS-V, IA, MCM, JME-S, JCA-D, AB-C, PP-V). PR-N and AG-M conducted DNA extractions, WES and all bioinformatics analyses. PR-N and JALE drafted the manuscript and all authors approved the final version of the manuscript.

Jose Antonio Lopez Escamez (JALE) is partially funded by INT18/00031 from ISCIII. This study was funded by the Luxembourg National Research Fund INTER/Mobility/17/11772209 Grant and EF-0247-2017 from Andalusian Health Government to JALE.

Authors declare no conflict of interest.

## Supplemental Digital Content

Supplemental Digital Content 1.docx

Supplemental Digital Content 2.docx

## Notes

**Conflicts of Interest and Source of Funding:** Jose Antonio Lopez Escamez (JALE) is partially funded by INT18/00031 from ISCIII. This study was funded by the Luxembourg National Research Fund INTER/Mobility/17/11772209 Grant and EF-0247-2017 from Andalusian Health Government to JALE. Authors declare no conflict of interest.

