## Supplemental Digital Content 1 for "Rare Variants in the *OTOG* Gene Are a Frequent Cause of Familial Meniere’s Disease"

**Table 1: Summary of the clinical information of familial MD patients carrying variants in *OTOG* gene.**

| Families | Code | | F1 | F2 | F3 | F4 | F5 | F6 | F7 | F8 | F9 |
| --- | --- | --- | --- | --- | --- | --- | --- | --- | --- | --- | --- |
|  | MD patients | Studied patient | II-4 | II-2 | II-2 | III-7 | II-2 | II-1 | II-3 | III-11 | III-1 |
|  |  | Other MD relatives | I-1; II-2 | I-2; II-1 | II-1 | II-1 | II-1; II-7; II-11 | II-2 | III-1 | II-6; III-7 | II-1; II-2 |
|  | Relatives with incomplete phenotype | Probable MD | II-1 | — | — | — | — | — | — | — | — |
|  |  | Episodic Vertigo | — | — | — | I-1; II-3; III-2 | — | — | — | II-1; II-9; III-13; III-15; III-19 | — |
|  |  | Hearing loss | — | — | — | — | — | — | — | II-3 | — |
| Clinical data of studied patient | Sex | | Female | Female | Female | Female | Female | Male | Female | Female | Female |
|  | Laterality | | Bilateral | Bilateral | Bilateral | Bilateral | Unilateral | Bilateral | Unilateral | Bilateral | Unilateral |
|  | Left ear | Low frequencies | Mild HL | Severe HL | Severe HL | Mild HL | MtoS HL | Moderate HL | MtoS HL | Mild HL | Severe HL |
|  |  | Mid frequencies | NH | Severe HL | Severe HL | Mild HL | MtoS HL | MtoS HL | MtoS HL | Mild HL | Severe HL |
|  |  | High frequencies | MtoS HL | Severe HL | Severe HL | Mild HL | MtoS HL | Profound HL | MtoS HL | Mild HL | Severe HL |
|  |  | Shape | Mild ski-slope | Flat | Flat | Flat | Flat | Ski-Slope | Flat | Flat | Flat |
|  | Right ear | Low frequencies | Severe HL | Moderate HL | Mild HL | MtoS HL | NH | Severe HL | NH | MtoS HL | NH |
|  |  | Mid frequencies | Moderate HL | Mild HL | Moderate HL | Mild HL | NH | Severe HL | NH | Severe HL | NH |
|  |  | High frequencies | Severe HL | Severe HL | Severe HL | Moderate HL | NH | Severe HL | NH | Profound HL | NH |
|  |  | Shape | Reverse-slope | Ski-Slope | Ski-Slope | Reverse-slope | NH | Flat | NH | Flat | NH |
|  | Other data | Age of onset | 50 | 31 | 52 | 30 | 53 | 31 | 51 | 41 | 33 |
|  |  | Headache | Paroxysmal hemicrania | MA | Tension headache | MO | No | Tension headache | No | No | Tension headache |
|  |  | Autoimmune disease | No | Hypothyroidism | No | No | No | No | No | No | No |

| Families | Code | | F10 | F11 | F12 | F13 | F14 |
| --- | --- | --- | --- | --- | --- | --- | --- |
|  | MD patients | Studied patient | II-2 | II-2 | III-6 | III-1 | II-3 |
|  |  | Other MD relatives | II-1 | II-1 | I-1; II-2 | I-1; III-2 | I-1; II-2 |
|  | Relatives with incomplete phenotype | Possible MD | — | — | — | — | II-1 |
|  |  | Vertigo | — | — | — | — | — |
|  |  | Hearing loss | — | — | — | II-3 | IV-4 |
| Clinical data of studied patient | Sex | | Female | Female | Male | Female | Male |
|  | Laterality | | Bilateral | Bilateral | Bilateral | Unilateral | Bilateral |
|  | Left ear | Low frequencies | MtoS HL | Severe HL | MtoS HL | NH | Moderate HL |
|  |  | Mid frequencies | Moderate HL | MtoS HL | MtoS HL | NH | Mild HL |
|  |  | High frequencies | MtoS HL | Profound HL | MtoS HL | NH | MtoS HL |
|  |  | Shape | Flat | Flat | Flat | NH | Reverse-slope |
|  | Right ear | Low frequencies | Profound HL | Mild HL | MtoS HL | Moderate HL | Mild HL |
|  |  | Mid frequencies | Profound HL | NH | MtoS HL | Moderate HL | Mild HL |
|  |  | High frequencies | Profound HL | Moderate HL | MtoS HL | Moderate HL | MtoS HL |
|  |  | Shape | Flat | Flat | Flat | Flat | Mild Ski-Slope |
|  | Other data | Age of onset | 53 | 38 | 42 | 42 | 40 |
|  |  | Headache | No | No | No | No | No |
|  |  | Autoimmune disease | Ulcerative colitis | No | No | No | No |

HL, hearing loss; MtoS, moderate-to-severe; MA, migraine with aura; MO, migraine without aura.

**Table 2: Clinical information of familial MD patients without variants in *OTOG* gene.**

| Code | Sex | Laterality | Ear affected | Age of onset | Relatives with MD | Headache | Autoimmune disease |
| --- | --- | --- | --- | --- | --- | --- | --- |
| F15 | Male | Unilateral | Left | 18 | Brother | Tension Headache | No |
| F16 | Female | Unilateral | Right | 34 | Grandmother | No | No |
| F17 | Male | Unilateral | Left | 33 | Sister and son | No | No |
| F18 | Female | Unilateral | Left | 50 | Uncle | No | — |
| F19 | Female | Bilateral | Both | 69 | Nephew | Tension Headache | No |
| F20 | Female | Bilateral | Both | 34 | Father | MA | No |
| F21 | Female | Bilateral | Both | 20 | Sister and brother | No | — |
| F22 | Female | Unilateral | Right | 20 | Mother and sister-in-law | MO | No |
| F23 | Female | Bilateral | Both | 16 | Mother | MA | Hypothyroidism |
| F24 | Female | Unilateral | Right | 42 | Mother | No | No |
| F25 | Female | Unilateral | Right | 31 | Mother and cousin | No | No |
| F26 | Male | Unilateral | Left | 54 | Father and brother | No | No |
| F27 | Male | Unilateral | Right | 25 | Mother | No | No |
| F28 | Female | Unilateral | Left | 19 | Mother | Tension headache | No |
| F29 | Female | Bilateral | Both | 40 | Daughter | Tension Headache | Type 1 Diabetes |
| F30 | Female | Unilateral | Right | 42 | Aunt | No | No |
| F31 | Female | Bilateral | Both | 28 | Mother | No | No |
| F32 | Male | Bilateral | Both | 31 | Daughter | Tension headache | No |
| F33 | Male | Bilateral | Both | 49 | Brother | No | No |
| F34 | Female | Unilateral | Right | 40 | Brother | No | No |
| F35 | Female | Bilateral | Both | 42 | Sister | No | No |
| F36 | Female | Bilateral | Both | 62 | Father | Tension headache | No |
| F37 | Female | Bilateral | Both | 46 | Twin sister | No | No |
| F38 | Male | Unilateral | Right | 70 | Sister | No | No |
| F39 | Female | Unilateral | Right | 56 | Father and Uncle | No | No |
| F40 | Female | Unilateral | Left | 54 | Sisters | Migraine | No |
| F41 | Male | Unilateral | Right | 42 | Mother | No | No |
| F42 | Female | Unilateral | Left | 29 | Mother | No | No |
| F43 | Female | Bilateral | Both | 14 | Father and Aunt | No | No |
| F44 | Female | Bilateral | Both | 15 | Father | No | No |
| F45 | Male | Bilateral | Both | 60 | Brother | MA | No |
| F46 | Female | Unilateral | Left | 20 | Mother | Yes | Arthritis |

MA, migraine with aura; MO, migraine without aura.

**Table 3: Clinical information of sporadic MD patients.**

| Code | Sex | Laterality | Ear affected | Age of onset | Headache | History of autoimmune disease |
| --- | --- | --- | --- | --- | --- | --- |
| S1 | Female | Unilateral | Right | 28 | Tension headache | No |
| S2 | Female | Unilateral | Right | 28 | Tension headache | Hypothyroidism |
| S3 | Male | Unilateral | Right | 33 | Tension headache | No |
| S4 | Male | Bilateral | Both | 16 | No | No |
| S5 | Female | Unilateral | Right | 27 | No | No |
| S6 | Female | Unilateral | Right | 20 | MA | Hypothyroidism |
| S7 | Female | Unilateral | Left | 14 | Tension headache | No |
| S8 | Male | Bilateral | Both | 35 | No | Psoriasis |
| S9 | Female | Unilateral | Left | 28 | No | No |
| S10 | Male | Bilateral | Both | 35 | No | Behçet syndrome |
| S11 | Female | Unilateral | Left | 31 | No | No |
| S12 | Female | Unilateral | Left | 30 | No | No |
| S13 | Female | Unilateral | Right | 27 | MO | No |
| S14 | Male | Bilateral | Both | 29 | No | No |
| S15 | Male | Bilateral | Both | 27 | No | Psoriasis |
| S16 | Female | Bilateral | Both | 33 | MO | Hughes syndrome |
| S17 | Female | Bilateral | Both | 24 | MA | No |
| S18 | Male | Unilateral | Left | 35 | No | No |
| S19 | Female | Unilateral | Left | 31 | MA | No |
| S20 | Female | Bilateral | Both | 23 | MO | Hypothyroidism |
| S21 | Female | Bilateral | Both | 35 | No | No |
| S22 | Female | Bilateral | Both | 22 | MO | Autoimmune arthralgia |
| S23 | Male | Unilateral | Left | 35 | No | No |
| S24 | Female | Unilateral | Left | 31 | No | No |
| S25 | Female | Unilateral | Right | 33 | No | No |
| S26 | Female | Unilateral | Right | 32 | No | No |
| S27 | Male | Bilateral | Both | 21 | Tension headache | No |
| S28 | Female | Bilateral | Both | 20 | No | No |
| S29 | Female | Bilateral | Both | 24 | No | No |
| S30 | Male | Bilateral | Both | 22 | Migraine | No |
| S31 | Female | Unilateral | Right | 21 | No | No |
| S32 | Female | Bilateral | Both | 27 | Tension headache | No |
| S33 | Male | Unilateral | Right | 30 | No | No |
| S34 | Male | Unilateral | Left | 25 | No | No |
| S35 | Female | Bilateral | Both | 25 | MA | No |
| S36 | Male | Bilateral | Both | 17 | No | Psoriasis |

MA, migraine with aura; MO, migraine without aura.

**Table 4: Sensorineural hearing loss genes selected for this study and information about its localizations, phenotypes and references.**

| Gene | Position | Phenotype | Reference |
| --- | --- | --- | --- |
| *ESPN* | chr1:6,484,848-6,521,430 | DFNB36 | <https://www.ncbi.nlm.nih.gov/pubmed/15930085> |
| *IFNLR1* | chr1:24,480,647-24,514,449 | DFNA2C | <https://www.ncbi.nlm.nih.gov/pubmed/29453195> |
| *GJB3* | chr1:35,246,790-35,251,970 | DFNA2B | <https://www.ncbi.nlm.nih.gov/pubmed/9843210> |
| *KCNQ4* | chr1:41,249,684-41,306,124 | DFNA2A | <https://www.ncbi.nlm.nih.gov/pubmed/10369879> |
| *BSND* | chr1:55,464,606-55,476,556 | DFNB73 | <https://www.ncbi.nlm.nih.gov/pubmed/19646679> |
| *ROR1* | chr1:64,239,690-64,647,181 | DFNB108 | <https://www.ncbi.nlm.nih.gov/pubmed/27162350> |
| *CDC14A* | chr1:100,810,584-100,985,833 | DFNB32 | <https://www.ncbi.nlm.nih.gov/pubmed/27259055> |
| COL11A1 | chr1:103,342,023-103,574,052 | DFNA37 | <https://www.ncbi.nlm.nih.gov/pubmed/30245514> |
| *GPSM2* | chr1:109,417,972-109,477,167 | DFNB82 | <https://www.ncbi.nlm.nih.gov/pubmed/20602914> |
| LMX1A | chr1:165,171,104-165,325,952 | DFNA7 | <https://www.ncbi.nlm.nih.gov/pubmed/29754270> |
| *NLRP3* | chr1:247,579,458-247,612,410 | DFNA34 | <https://www.ncbi.nlm.nih.gov/pubmed/28847925> |
| *OTOF* | chr2:26,680,071-26,781,566 | DFNB9 | <https://www.ncbi.nlm.nih.gov/pubmed/10903124> |
| *PNPT1* | chr2:55,861,198-55,921,045 | DFNB70 | <https://www.ncbi.nlm.nih.gov/pubmed/23084290> |
| *ELMOD3* | chr2:85,581,517-85,618,875 | DFNB88 | <https://www.ncbi.nlm.nih.gov/pubmed/24039609> |
| *PJVK* | chr2:179,316,163-179,326,117 | DFNB59 | <https://www.ncbi.nlm.nih.gov/pubmed/16804542> |
| *TMIE* | chr3:46,742,823-46,752,413 | DFNB6 | <https://www.ncbi.nlm.nih.gov/pubmed/12145746> |
| *ILDR1* | chr3:121,706,170-121,741,127 | DFNB42 | <https://www.ncbi.nlm.nih.gov/pubmed/21255762> |
| *MCM2* | chr3:127,317,066-127,341,279 | DFNA70 | <https://www.ncbi.nlm.nih.gov/pubmed/26196677> |
| *CCDC50* | chr3:191,046,866-191,116,459 | DFNA344 | <https://www.ncbi.nlm.nih.gov/pubmed/17503326> |
| *WFS1* | chr4:6,271,576-6,304,992 | DFNA6 | <https://www.ncbi.nlm.nih.gov/pubmed/10424813> |
| *GRXCR1* | chr4:42,895,283-43,032,675 | DFNB25 | <https://www.ncbi.nlm.nih.gov/pubmed/20137778> |
| REST | chr4:57,774,042-57,802,010 | DFNA27 | <https://www.ncbi.nlm.nih.gov/pubmed/29961578> |
| GAB1 | chr4:144,257,915-144,395,721 | DFNB26 | <https://www.ncbi.nlm.nih.gov/pubmed/29408807> |
| *MARVELD2* | chr5:68,710,939-68,740,157 | DFNB49 | <https://www.ncbi.nlm.nih.gov/pubmed/15538632> |
| *BDP1* | chr5:70,751,442-70,863,649 | DFNB112 | <https://www.ncbi.nlm.nih.gov/pubmed/24312468> |
| *PPIP5K2* | chr5:102,455,853-102,548,500 | DFNB100 | <https://www.ncbi.nlm.nih.gov/pubmed/15538632> |
| *SLC22A4* | chr5:131,630,136-131,679,899 | DFNB60 | <https://www.ncbi.nlm.nih.gov/pubmed/27023905> |
| Gene | Position | Phenotype | Reference |
| *DIAPH1* | chr5:140,894,583-140,998,622 | DFNA1 | <https://www.ncbi.nlm.nih.gov/pubmed/9360932> |
| *GRXCR2* | chr5:145,239,296-145,252,531 | DFNB101 | <https://www.ncbi.nlm.nih.gov/pubmed/24619944> |
| *POU4F3* | chr5:145,718,587-145,720,083 | DFNA15 | <https://www.ncbi.nlm.nih.gov/pubmed/18228599> |
| *SERPINB6* | chr6:2,948,393-2,972,399 | DFNB91 | <https://www.ncbi.nlm.nih.gov/pubmed/20451170> |
| *DCDC2* | chr6:24,171,983-24,383,520 | DFNB66 | <https://www.ncbi.nlm.nih.gov/pubmed/25601850> |
| *RIPOR2* | chr6:24,797,601-25,042,238 | DFNB104 | <https://www.ncbi.nlm.nih.gov/pubmed/24958875> |
| COL11A2 | chr6:33,130,458-33,160,276 | DFNA13 | <https://www.ncbi.nlm.nih.gov/pubmed/10581026> |
| *LHFPL5* | chr6:35,773,070-35,801,651 | DFNB67 | <https://www.ncbi.nlm.nih.gov/pubmed/16752389> |
| *CLIC5* | chr6:45,866,188-46,048,132 | DFNB103 | <https://www.ncbi.nlm.nih.gov/pubmed/17021174> |
| *MYO6* | chr6:76,458,909-76,629,254 | DFNA22 | <https://www.ncbi.nlm.nih.gov/pubmed/11468689> |
| *CD164* | chr6:109,687,717-109,703,762 | DFNA66 | <https://www.ncbi.nlm.nih.gov/pubmed/26197441> |
| *EYA4* | chr6:133,561,736-133,853,258 | DFNA10 | <https://www.ncbi.nlm.nih.gov/pubmed/17567890> |
| GSDME | chr7:24,737,972-24,809,244 | DFNA5 | <https://www.ncbi.nlm.nih.gov/pubmed/9771715> |
| PDE1C | chr7:31,790,793-32,338,941 | DFNA74 | <https://www.ncbi.nlm.nih.gov/pubmed/29860631> |
| *ADCY1* | chr7:45,613,739-45,762,715 | DFNB44 | <https://www.ncbi.nlm.nih.gov/pubmed/15583425> |
| *HGF* | chr7:81,328,322-81,399,754 | DFNB39 | <https://www.ncbi.nlm.nih.gov/pubmed/19576567> |
| TRRAP | chr7:98,475,556-98,610,866 | DFNA | <https://www.ncbi.nlm.nih.gov/pubmed/31231791> |
| *SLC26A5* | chr7:102,993,177-103,086,624 | DFNB61 | <https://www.ncbi.nlm.nih.gov/pubmed/12719379> |
| *SLC26A4* | chr7:107,301,080-107,358,254 | DFNB4 | <https://www.ncbi.nlm.nih.gov/pubmed/16570074> |
| *MET* | chr7:116,312,444-116,438,440 | DFNB97 | <https://www.ncbi.nlm.nih.gov/pubmed/25941349> |
| MIRN96 | chr7:129,414,532-129,414,609 | DFNA50 | <https://www.ncbi.nlm.nih.gov/pubmed/19363479> |
| *ESRP1* | chr8:95,653,302-95,719,694 | DFNB109 | <https://www.ncbi.nlm.nih.gov/pubmed/29107558> |
| *GRHL2* | chr8:102,504,660-102,681,954 | DFNA28 | <https://www.ncbi.nlm.nih.gov/pubmed/12393799> |
| *TJP2* | chr9:71,736,209-71,870,124 | DFNA51 | <https://www.ncbi.nlm.nih.gov/pubmed/20602916> |
| *TMC1* | chr9:75,136,717-75,451,267 | DFNB7 | <https://www.ncbi.nlm.nih.gov/pubmed/18616530> |
| *WHRN* | chr9:117,164,360-117,267,736 | DFNB31 | <https://www.ncbi.nlm.nih.gov/pubmed/12833159> |
| *TNC* | chr9:117,782,805-117,880,536 | DFNA56 | <https://www.ncbi.nlm.nih.gov/pubmed/23936043> |
| *TPRN* | chr9:140,086,069-140,098,645 | DFNB79 | <https://www.ncbi.nlm.nih.gov/pubmed/20170898> |
| *MYO3A* | chr10:26,223,002-26,501,465 | DFNB30 | <https://www.ncbi.nlm.nih.gov/pubmed/12032315> |
| Gene | Position | Phenotype | Reference |
| *PCDH15* | chr10:55,562,531-57,387,702 | DFNB23 | <https://www.ncbi.nlm.nih.gov/pubmed/14570705> |
| *CDH23* | chr10:73,156,691-73,575,704 | DFNB12 | <https://www.ncbi.nlm.nih.gov/pubmed/11090341> |
| *C10orf105* | chr10:73,471,458-73,497,581 | DFNB12 | <https://www.ncbi.nlm.nih.gov/pubmed/11090341> |
| *PDZD7* | chr10:102,767,440-102,790,914 | DFNB57 | <https://www.ncbi.nlm.nih.gov/pubmed/26849169> |
| *EPS8L2* | chr11:694,438-727,727 | DFNB106 | <https://www.ncbi.nlm.nih.gov/pubmed/26282398> |
| *USH1C* | chr11:17,515,442-17,565,963 | DFNB18A | <https://www.ncbi.nlm.nih.gov/pubmed/12107438> |
| *OTOG* | chr11:17,568,920-17,668,697 | DFNB18B | <https://www.ncbi.nlm.nih.gov/pubmed/23122587> |
| *CABP2* | chr11:67,286,383-67,290,899 | DFNB93 | <https://www.ncbi.nlm.nih.gov/pubmed/22981119> |
| *LRTOMT* | chr11:71,791,377-71,821,828 | DFNB63 | <https://www.ncbi.nlm.nih.gov/pubmed/18953341> |
| *ANAPC15* | chr11:71,817,424-71,823,826 | DFNB63 | <https://www.ncbi.nlm.nih.gov/pubmed/18953341> |
| *MYO7A* | chr11:76,839,310-76,926,286 | DFNA11A | <https://www.ncbi.nlm.nih.gov/pubmed/8776602> |
| *NARS2* | chr11:78,147,007-78,285,919 | DFNB94 | <https://www.ncbi.nlm.nih.gov/pubmed/25807530> |
| *RDX* | chr11:110,045,605-110,167,447 | DFNB24 | <https://www.ncbi.nlm.nih.gov/pubmed/17226784> |
| MPZL2 | chr11:118,124,118-118,135,251 | DFNB111 | <https://www.ncbi.nlm.nih.gov/pubmed/29961571> |
| *TECTA* | chr11:120,971,882-121,062,202 | DFNA8 | <https://www.ncbi.nlm.nih.gov/pubmed/16718611> |
| *EPS8* | chr12:15,773,075-16,035,263 | DFNB102 | <https://www.ncbi.nlm.nih.gov/pubmed/24741995> |
| *MSRB3* | chr12:65,672,423-65,882,024 | DFNB74 | <https://www.ncbi.nlm.nih.gov/pubmed/21185009> |
| *OTOGL* | chr12:80,603,233-80,772,870 | DFNB84B | <https://www.ncbi.nlm.nih.gov/pubmed/23122586> |
| *PTPRQ* | chr12:80,799,774-81,073,968 | DFNB84 | <https://www.ncbi.nlm.nih.gov/pubmed/20346435> |
| *KITLG* | chr12:88,886,570-88,974,628 | DFNA69 | <https://www.ncbi.nlm.nih.gov/pubmed/26522471> |
| *SLC17A8* | chr12:100,750,857-100,815,837 | DFNA25 | <https://www.ncbi.nlm.nih.gov/pubmed/11115382> |
| *DIABLO* | chr12:122,692,209-122,712,081 | DFNA64 | <https://www.ncbi.nlm.nih.gov/pubmed/21722859> |
| *P2RX2* | chr12:133,195,366-133,198,972 | DFNA41 | <https://www.ncbi.nlm.nih.gov/pubmed/24211385> |
| *GJB2* | chr13:20,761,602-20,767,114 | DFNA3A | <https://www.ncbi.nlm.nih.gov/pubmed/9620796> |
| *GJB6* | chr13:20,796,101-20,806,534 | DFNA3B | <https://www.ncbi.nlm.nih.gov/pubmed/10471490> |
| *COCH* | chr14:31,343,720-31,364,271 | DFNA9 | <https://www.ncbi.nlm.nih.gov/pubmed/9806553> |
| *SIX1* | chr14:61,110,133-61,124,977 | DFNA23 | <https://www.ncbi.nlm.nih.gov/pubmed/15141091> |
| *ESRRB* | chr14:76,776,957-76,968,180 | DFNB35 | <https://www.ncbi.nlm.nih.gov/pubmed/18179891> |
| *STRC* | chr15:43,891,596-44,010,458 | DFNB16 | <https://www.ncbi.nlm.nih.gov/pubmed/11687802> |
| Gene | Position | Phenotype | Reference |
| *DMXL2* | chr15:51,739,908-51,915,030 | DFNA71 | <https://www.ncbi.nlm.nih.gov/pubmed/27657680> |
| *CIB2* | chr15:78,396,948-78,423,886 | DFNB48 | <https://www.ncbi.nlm.nih.gov/pubmed/23023331> |
| *HOMER2* | chr15:83,509,838-83,654,661 | DFNA68 | <https://www.ncbi.nlm.nih.gov/pubmed/25816005> |
| *TBC1D24* | chr16:2,525,147-2,555,735 | DFNB86 | <https://www.ncbi.nlm.nih.gov/pubmed/24387994> |
| CLDN9 | chr16:3,062,457-3,064,506 | DFNB | <https://www.ncbi.nlm.nih.gov/pubmed/31175426> |
| CRYM | chr16:21,250,195-21,314,404 | DFNA40 | <https://www.ncbi.nlm.nih.gov/pubmed/12471561> |
| *OTOA* | chr16:21,689,835-21,772,050 | DFNB22 | <https://www.ncbi.nlm.nih.gov/pubmed/11972037> |
| *KARS* | chr16:75,661,622-75,682,541 | DFNB89 | <https://www.ncbi.nlm.nih.gov/pubmed/23768514> |
| SPNS2 | chr17:4,402,129-4,443,228 | DFNB115 | <https://www.ncbi.nlm.nih.gov/pubmed/30973865> |
| *MYO15A* | chr17:18,012,020-18,083,116 | DFNB3 | <https://www.ncbi.nlm.nih.gov/pubmed/9603736> |
| GRAP | chr17:18,923,986-18,950,950 | DFNB114 | <https://www.ncbi.nlm.nih.gov/pubmed/30610177> |
| *TMEM132E* | chr17:32,907,768-32,966,337 | DFNB99 | <https://www.ncbi.nlm.nih.gov/pubmed/25331638> |
| *WBP2* | chr17:73,841,780-73,852,588 | DFNB107 | <https://www.ncbi.nlm.nih.gov/pubmed/26881968> |
| *ACTG1* | chr17:79,476,997-79,490,873 | DFNA20 | <https://www.ncbi.nlm.nih.gov/pubmed/14684684> |
| *LOXHD1* | chr18:44,056,935-44,236,996 | DFNB77 | <https://www.ncbi.nlm.nih.gov/pubmed/21465660> |
| *GIPC3* | chr19:3,585,551-3,593,539 | DFNB15 | <https://www.ncbi.nlm.nih.gov/pubmed/21660509> |
| *S1PR2* | chr19:10,332,109-10,341,948 | DFNB68 | <https://www.ncbi.nlm.nih.gov/pubmed/26805784> |
| *SYNE4* | chr19:36,494,002-36,499,695 | DFNB76 | <https://www.ncbi.nlm.nih.gov/pubmed/23348741> |
| *CEACAM16* | chr19:45,202,421-45,213,986 | DFNA4B | <https://www.ncbi.nlm.nih.gov/pubmed/25589040> |
| *MYH14* | chr19:50,691,443-50,813,802 | DFNA4A | <https://www.ncbi.nlm.nih.gov/pubmed/15015131> |
| *OSBPL2* | chr20:60,813,580-60,871,269 | DNFA67 | <https://www.ncbi.nlm.nih.gov/pubmed/25759012> |
| *CLDN14* | chr21:37,832,919-37,948,867 | DFNB29 | <https://www.ncbi.nlm.nih.gov/pubmed/11163249> |
| *TMPRSS3* | chr21:43,791,996-43,816,955 | DFNB8 | <https://www.ncbi.nlm.nih.gov/pubmed/11907649> |
| *TSPEAR* | chr21:45,917,775-46,131,495 | DFNB98 | <https://www.ncbi.nlm.nih.gov/pubmed/22678063> |
| *MYH9* | chr22:36,677,323-36,784,063 | DFNA17 | <https://www.ncbi.nlm.nih.gov/pubmed/11023810> |
| *TRIOBP* | chr22:38,092,995-38,172,563 | DFNB28 | <https://www.ncbi.nlm.nih.gov/pubmed/16385458> |
| *SMPX* | chrX:21,724,090-21,776,281 | DFNX4 | <https://www.ncbi.nlm.nih.gov/pubmed/21549342> |
| *POU3F4* | chrX:82,763,269-82,764,775 | DFNX2 | <https://www.ncbi.nlm.nih.gov/pubmed/7839145> |
| *PRPS1* | chrX:106,871,654-106,894,256 | DFNX1 | <https://www.ncbi.nlm.nih.gov/pubmed/20021999> |
| Gene | Position | Phenotype | Reference |
| *COL4A6* | chrX:107,386,780-107,682,727 | DFNX6 | <https://www.ncbi.nlm.nih.gov/pubmed/23714752> |
| *AIFM1* | chrX:129,263,337-129,299,861 | DFNX5 | <https://www.ncbi.nlm.nih.gov/pubmed/25986071> |

**Table 5: Rare variants found in the SRVA for familial MD cases.**

| Variant | Gene | Exon | Fam MD Code | MAF NFE | MAF CSVS | CADD | ACMG |
| --- | --- | --- | --- | --- | --- | --- | --- |
| chr1:6488328C>T | *ESPN* | 2 | F31 | 0.00040 | 0.0024 | 35 | Uncertain Significance |
| chr1:35250892T>G | *GJB3* | 2 | F32 | 0.00010 | 0.0003 | 24.5 | Likely benign |
| chr1:109472327C>T | *GPSM2* | 15 | F19 | 0.000015 | 0.00030 | 34 | Benign |
| chr2:26696027G>C | *OTOF* | 29 | F19 | 0.00090 | 0.0012 | 22.7 | Uncertain Significance |
| chr2:26741960C>T | *OTOF* | 4 | F3 | 0.000088 | 0.00091 | 26.2 | Likely benign |
| chr3:46747377C>T | *TMIE* | 2 | F45 | 0.00040 | 0.00030 | 24.2 | Uncertain Significance |
| chr3:127325493G>C | *MCM2* | 6 | F26 | 0.00070 | 0.00030 | 27.8 | Uncertain Significance |
| chr5:70793116A>C | *BDP1* | 13 | F42 | 0.000030 | 0.00030 | 23.9 | Uncertain Significance |
| chr5:102530663C>T | *PPIP5K2* | 30 | F27 | 0.00050 | 0.00061 | 23.7 | Uncertain Significance |
| chr5:140953193G>C | *DIAPH1* | 16 | F39 | 0.00040 | 0.00061 | 18.90 | Uncertain Significance |
| chr6:2954942G>T | *SERPINB6* | 5 | F1 | 0.00080 | 0.0024 | 24.7 | Uncertain Significance |
| chr7:107329557T>C | *SLC26A4* | 9 | F27 | 0.00060 | 0.0043 | 31 | Likely benign |
| chr7:107350627G>A | *SLC26A4* | 19 | F39 | 0.00020 | 0.0015 | 23.2 | Benign |
| chr7:116411646C>T | *MET* | 13 | F41 | 0.000060 | 0.00030 | 21.0 | Uncertain Significance |
| chr9:71845053G>T | *TJP2* | 12 | F19 | 0.000045 | 0.00030 | 32 | Uncertain Significance |
| chr10:55581760C>T | *PCDH15* | 35 | F28 | 0.00010 | 0.0012 | 15.06 | Uncertain Significance |
| chr10:55912942C>T | *PCDH15* | 15 | F39 | 0.00020 | 0.0015 | 24.6 | benign |
| chr10:56106198T>C | *PCDH15* | 7 | F24 | 0.000029 | 0.0018 | 22.7 | Uncertain Significance |
| chr10:73553197G>A | *CDH23* | 46 | F35 | 0 | 0.00030 | 17.72 | Uncertain Significance |
| chr11:720123G>A | *EPS8L2* | 5 | F43 | 0.000061 | 0.00030 | 32 | Uncertain Significance |
| chr11:17531093G>C | *USH1C* | 18 | F27 | 0.00080 | 0.00030 | 24.0 | Uncertain Significance |
| chr11:17574758G>A | *OTOG* | 5 | F1; F14 | 0.00080 | 0.0034 | 24.8 | Uncertain Significance |
| chr11:67287311A>G | *CABP2* | 6 | F44 | 0.00060 | 0.00030 | 26.1 | Uncertain Significance |
| chr11:76885923G>A | *MYO7A* | 17 | F37 | 0.00010 | 0.0030 | 34 | Uncertain Significance |
| chr11:76922875G>A | *MYO7A* | 46 | F27 | 0.00040 | 0.00030 | 22.1 | Uncertain Significance |
| chr11:121028725T>C | *TECTA* | 13 | F8 | 0.000091 | 0.00091 | 22.3 | Uncertain Significance |
| chr12:15823797C>T | *EPS8* | 4 | F28 | 0.00080 | 0.0024 | 18.82 | Uncertain Significance |
| chr12:80752642T>G | *OTOGL* | 51 | F31 | 0 | 0.00030 | 27.6 | Uncertain Significance |
| chr12:100751192C>T | *SLC17A8* | 1 | F33 | 0.00030 | 0.0012 | 19.98 | Benign |
| chr14:31355389A>G | *COCH* | 11 | F23 | 0.0010 | 0.0021 | 22.0 | Likely benign |
| Variant | Gene | Exon | Fam MD Code | MAF NFE | MAF CSVS | CADD | ACMG |
| chr16:21739665C>T | *OTOA* | 19 | F25 | 0.00020 | 0.0040 | 19.35 | Likely benign |
| chr18:44109147C>T | *LOXHD1* | 29 | F31 | 0.00060 | 0.0012 | 19.82 | Likely benign |
| chr18:44184084C>T | *LOXHD1* | 7 | F27 | 0.00030 | 0.0024 | 26.8 | Uncertain Significance |
| chr19:3586840G>A | *GIPC3* | 3 | F24 | 0.00040 | 0.00030 | 23.4 | Likely benign |
| chr19:50785088A>G | *MYH14* | 33 | F4 | 0 | 0.00030 | 31 | Uncertain Significance |
| chr19:50810310C>T | *MYH14* | 41 | F8 | 0 | 0.00061 | 34 | Uncertain Significance |
| chr21:43796787C>T | *TMPRSS3* | 11 | F17 | 0.000030 | 0.00030 | 18.18 | Uncertain Significance |
| chr21:45945541C>T | *TSPEAR* | 9 | F16 | 0.000030 | 0.00030 | 34 | Uncertain Significance |
| chr22:36697620G>A | *MYH9* | 21 | F45 | 0.00060 | 0.00030 | 23.1 | Uncertain Significance |
| chr22:38165350C>T | *TRIOBP* | 21 | F45 | 0.000021 | 0.00030 | 33 | Uncertain Significance |

MAF NFE, minor allele frequency in the Non-Finnish European population from ExAC; MAF CSVS, minor allele frequency in the Collaborative Spanish Variant Server; CADD, Combined Annotation Dependent Depletion Score; ACMG, pathogenicity according to assessed according to the guidelines described by the American College of Medical Genetics and Genomics and the Association for Molecular Pathology.

**Table 6: Rare variants found in the SRVA for sporadic MD cases.**

| Variant | Gene | Exon | Sporadic cases | MAF NFE | MAF CSVS | CADD |
| --- | --- | --- | --- | --- | --- | --- |
| chr1:24484051C>T | *IFNLR1* | 7 | S33 | 0.000045 | 0 | 20.6 |
| chr1:247588214G>A | *NLRP3* | 5 | S26 | 0.0010 | 0.0010 | 22.3 |
| chr2:55863449C>T | *PNPT1* | 28 | S2 | 0.000045 | — | 23.9 |
| chr2:179319145G>A | *DFNB59* | 3 | S32 | 0.00040 | — | 25.7 |
| chr4:6279258C>T | *WFS1* | 2 | S10 | 0 | — | 35 |
| chr4:6303833G>A | *WFS1* | 8 | S17 | 0 | — | 22.8 |
| chr5:70785335G>C | *BDP1* | 10 | S24 | 0.00020 | 0.0010 | 16.52 |
| chr5:70806540A>G | *BDP1* | 17 | S4 | 0.000030 | — | 21.7 |
| chr5:70855826C>G | *BDP1* | 37 | S24 | 0.00060 | 0.0010 | 21.9 |
| chr5:145719938C>G | *POU4F3* | 2 | S9 | 0.00050 | 0 | 26.4 |
| chr7:45632443C>T | *ADCY1* | 3 | S11 | 0.00050 | — | 23.6 |
| chr7:103018088T>G | *SLC26A5* | 18 | S4 | 0.00020 | 0.0030 | 25.3 |
| chr9:117819627C>T | *TNC* | 15 | S23 | 0.000075 | 0 | 17.16 |
| chr9:117852972C>T | *TNC* | 2 | S22 | 0 | — | 18.08 |
| chr9:140093585G>A | *TPRN* | 1 | S6 | 0.00040 | 0.0010 | 24.3 |
| chr10:73406348G>A | *CDH23* | 13 | S22 | 0.00010 | 0.0010 | 24.8 |
| chr10:73466770G>A | *CDH23* | 25 | S18 | 0.00010 | — | 24.7 |
| chr10:102775470G>A | *PDZD7* | 11 | S27 | 0.00040 | — | 20.6 |
| chr10:102783196G>A | *PDZD7* | 4 | S3 | 0.00020 | 0 | 32 |
| chr11:17522618C>T | *USH1C* | 23 | S29 | 0 | — | 26.8 |
| chr11:17615604C>T | *OTOG* | 28 | S12 | 0.00090 | 0.0070 | 23.1 |
| chr11:17615655C>T | *OTOG* | 28 | S12 | 0.00090 | 0.0070 | 33 |
| chr11:17632279C>T | *OTOG* | 36 | S1; S23 | 0 | — | 26.4 |
| chr11:17653443C>T | *OTOG* | 41 | S13 | 0.00060 | 0.0030 | 34 |
| chr11:76883864G>A | *MYO7A* | 16 | S26 | 0.00030 | 0.0010 | 34 |
| chr11:76922974G>A | *MYO7A* | 46 | S31 | 0.000062 | — | 26.9 |
| chr11:78239889C>G | *NARS2* | 6 | S31 | 0.00020 | 0 | 22.7 |
| chr11:120998763G>A | *TECTA* | 8 | S18 | 0.000075 | — | 24.9 |
| chr12:80752642T>G | *OTOGL* | 51 | S27 | 0 | 0 | 27.6 |
| chr12:80771692G>A | *OTOGL* | 58 | S26 | 0.00020 | 0.0010 | 24.3 |
| chr13:20797249T>C | *GJB6* | 5 | S36 | 0 | — | 18.08 |
| chr15:83532948T>C | *HOMER2* | 4 | S25 | 0.00040 | 0 | 23.3 |
| chr16:2546934C>T | *TBC1D24* | 2 | S27 | 0.00030 | 0.0010 | 16.76 |
| chr16:75663344C>T | *KARS* | 13 | S4 | 0.000029 | — | 23.6 |
| chr17:18039039G>T | *MYO15A* | 12 | S5 | 0.0010 | 0.0030 | 22.9 |
| chr17:18049397C>T | *MYO15A* | 29 | S35 | 0.000046 | 0 | 24.3 |
| chr17:73845743G>C | *WBP2* | 3 | S35 | 0.000014 | — | 25.4 |
| chr22:36714278C>T | *MYH9* | 11 | S1 | 0.000030 | — | 33 |
| chr22:38130521C>T | *TRIOBP* | 9 | S3 | 0.000074 | 0.0010 | 20.6 |
| chr22:38164179C>T | *TRIOBP* | 19 | S18 | 0.000060 | 0.0020 | 26.2 |
| chr22:38167706T>C | TRIOBP | 22 | S3 | 0.000030 | 0.0010 | 23.6 |

MAF NFE, minor allele frequency in the Non-Finnish European population from ExAC; MAF CSVS, minor allele frequency in the Collaborative Spanish Variant Server; CADD, Combined Annotation Dependent Depletion Score.

| Position | Exon | Sporadic cases | MAF NFE | MAF CSVS | CADD | Domain |
| --- | --- | --- | --- | --- | --- | --- |
| chr11:17578774G>A | 7 | S24 | 0.0090 | 0.018 | 15.95 | vWD |
| chr11:17591922C>T | 16 | S1; S12 | 0.032 | 0.030 | 25.4 | vWD |
| chr11:17615604C>T | 27 | S12 | 0.00090 | 0.0070 | 23.1 | vWD |
| chr11:17615655C>T | 27 | S12 | 0.00090 | 0.0070 | 33 | vWD |
| chr11:17631453C>T | 35 | S11; S24 | 0.016 | 0.016 | 12.89 | — |
| chr11:17632099G>A | 35 | S5 | 0.00060 | 0.0030 | 0.073 | — |
| chr11:17632279C>T | 35 | S1; S23 | 0 | — | 26.4 | — |
| chr11:17653443C>T | 40 | S13 | 0.00060 | 0.0030 | 34 | C8 |
| chr11:17655748G>A | 43 | S1 | 0.031 | 0.022 | 14.23 | — |
| chr11:17656672G>A | 45 | S9 | 0.0034 | 0.0040 | 31 | — |
| chr11:17663747G>A | 52 | S7 | 0.0058 | 0.0060 | 19.41 | — |
| chr11:17667139G>C | 54 | S12; S20 | 0.018 | 0.019 | 27.2 | CT |

**Table 7: Rare variants found in the gene burden analysis in *OTOG* gene for sporadic MD cases.**

MAF NFE, minor allele frequency in the Non-Finnish European population from ExAC; MAF CSVS, minor allele frequency in the Collaborative Spanish Variant Server; CADD, Combined Annotation Dependent Depletion Score; vWD, von Willebrand factor type D domain; CT, cysteine knot domain.
