## Supplemental Digital Content 1 for "Rare Variants in the *OTOG* Gene Are a Frequent Cause of Familial Meniere’s Disease"

**Figure 1: Pedigrees of the 14 families carrying variants in the *OTOG* gene. Seven families showed incomplete penetrance (1A) and seven families showed complete penetrance (1B)**.

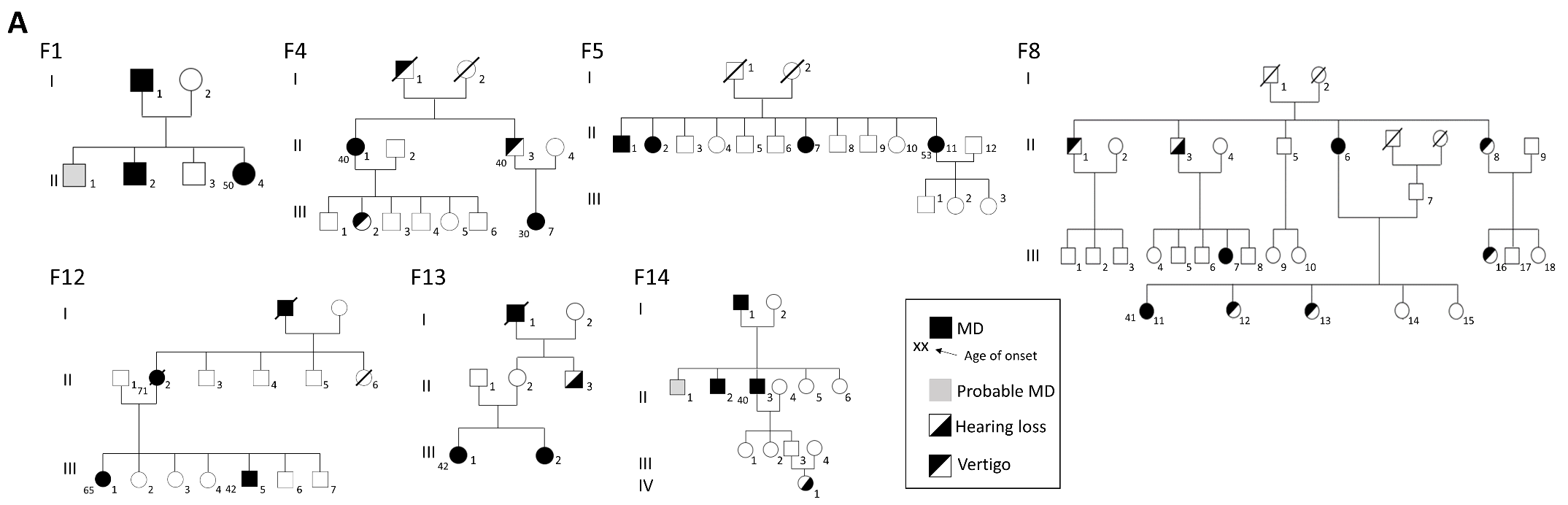

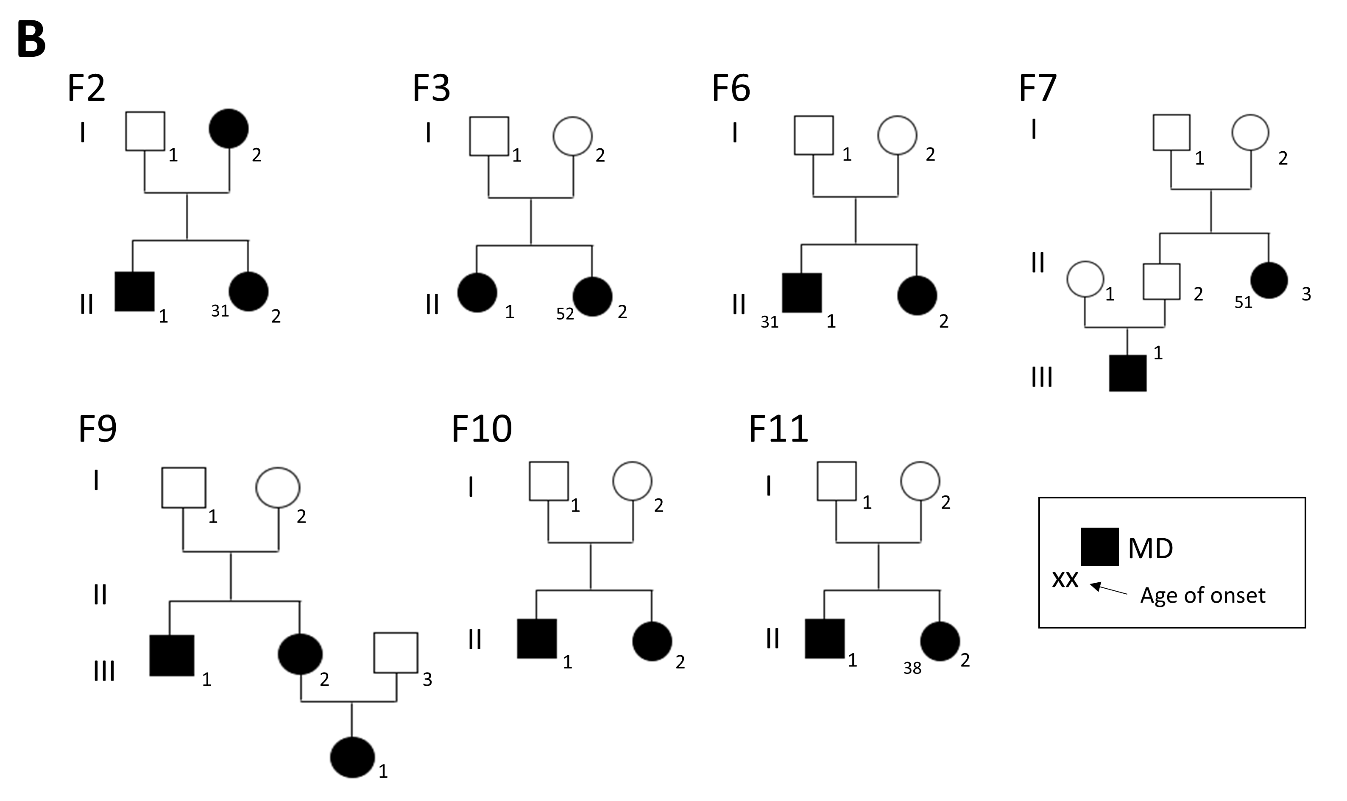

In total, 38 individuals with MD where diagnosed in these families. Likewise, 5 families (F1, F4, F8, F13 and F14) showed partial syndromes (probable MD, hearing loss or vertigo), suggesting phenotypic heterogeneity. Furthermore, incomplete penetrance was observed in 7 families (F1, F4, F5, F8, F12, F13 and F14).

**Figure 2: Serial pure tone audiograms for the 14 familial MD patients with variants in *OTOG* gene**.

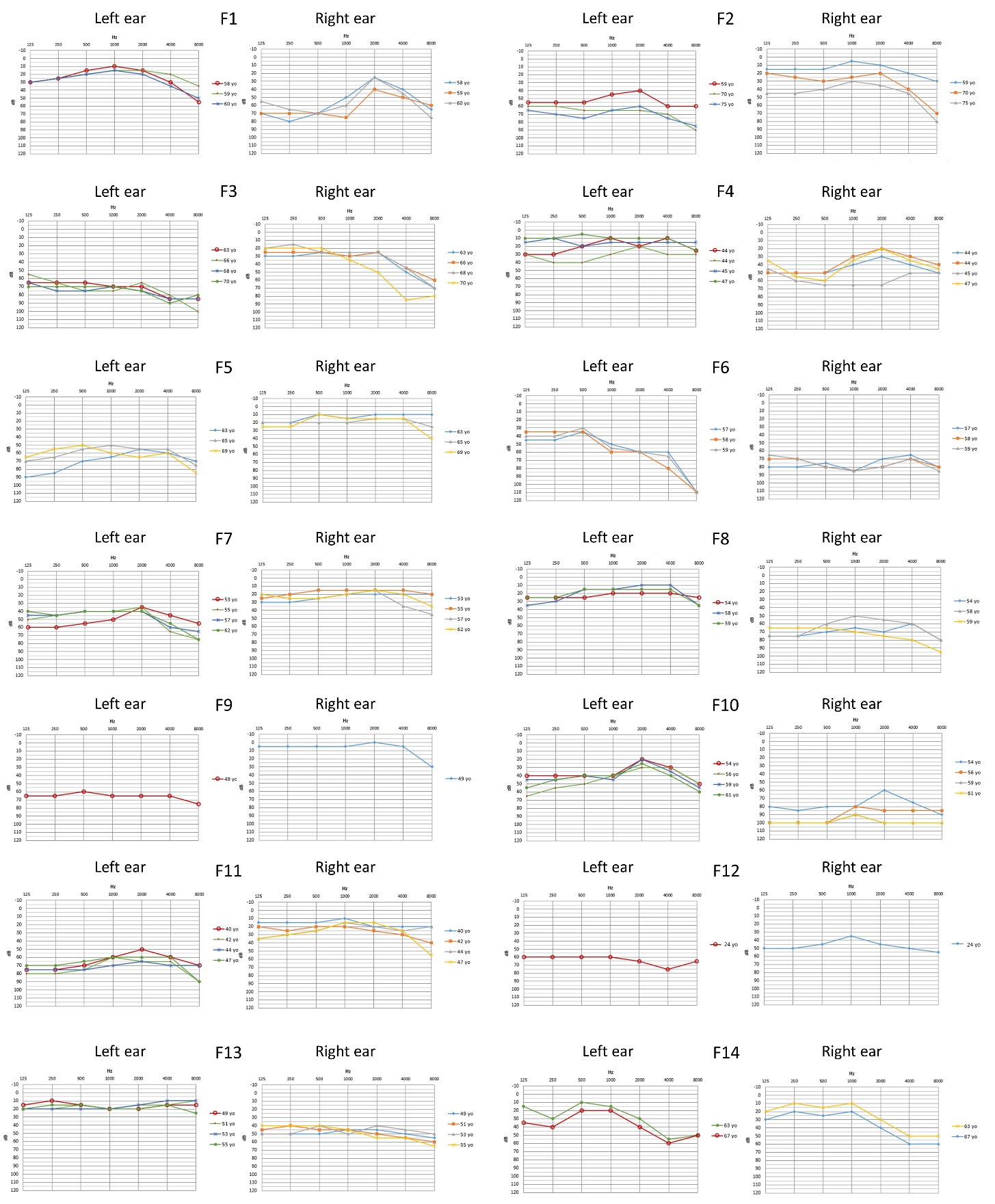
 yo, years old.

**Figure 3: Flowchart summarizing the bioinformatic analysis on sporadic MD cases**.

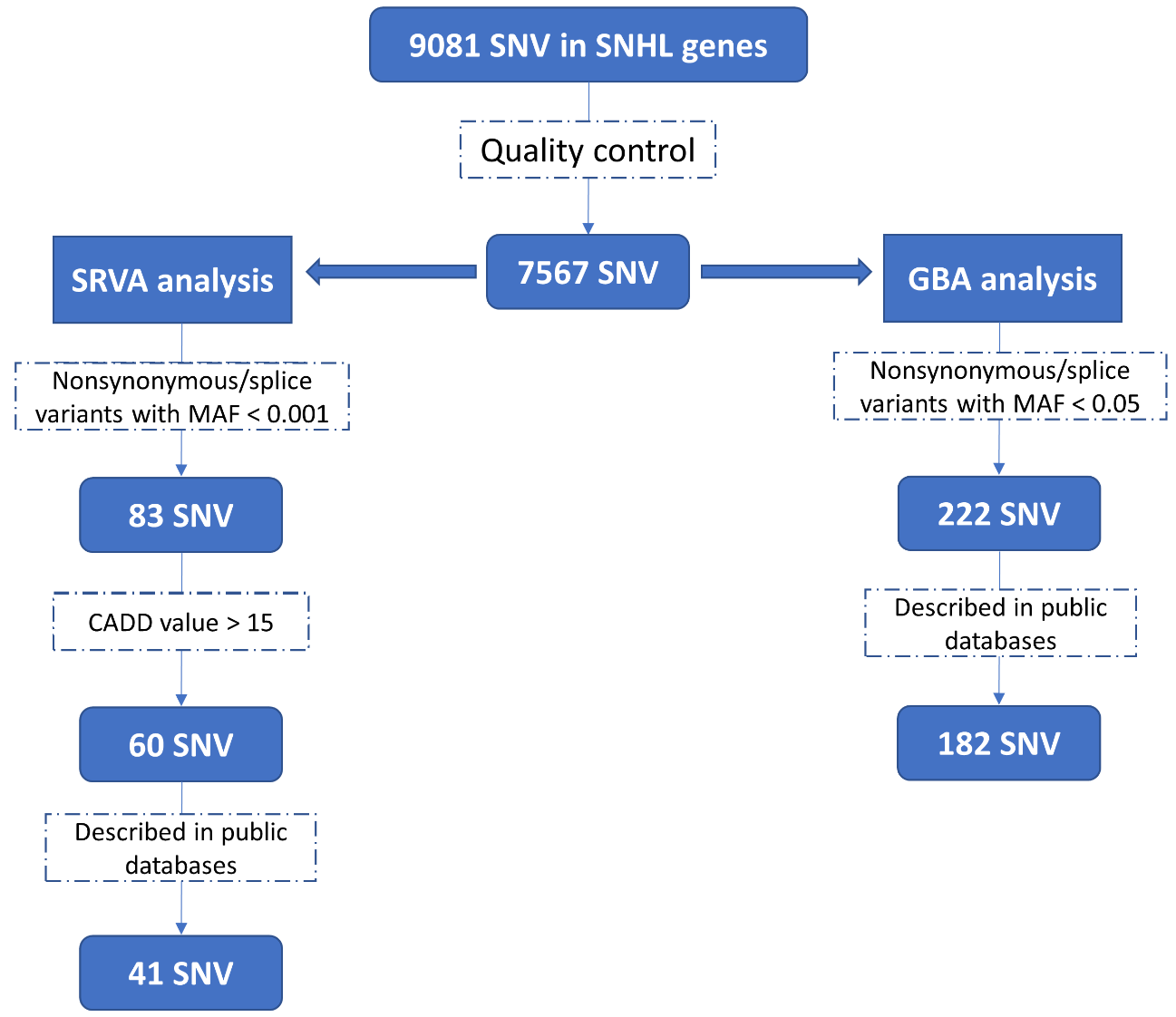

On the left, single rare variant analysis (SRVA) and prioritization pipeline. On the right, the gene burden analysis (GBA) pipeline. SNV, single nucleotide variants; CADD, Combined Annotation Dependent Depletion Score.
